# A New Species of Seedeater (Emberizidae: *Sporophila*) from the Iberá Grasslands, in Northeast Argentina

**DOI:** 10.1101/046318

**Authors:** Adrián S. Di Giacomo, Bernabé López-Lanús, Cecilia Kopuchian

**Author notes:** Corresponding autor.

## Abstract

We describe a new species of capuchino of the genus *Sporophila* (Emberizidae) of the Esteros del Iberá, province of Corrientes, in northeastern Argentina. This species would have remained unidentified due to lack of ornithological explorations in the central area of the Esteros del Iberá. It has been confused with immature individuals of other *Sporophila* species. We made observations of behavior and habitat, playback experiments, comparative analyzes of the vocalizations and plumage with other sympatric species of the same genus, and we found this species, which we have named *Sporophila iberaensis*, inhabiting wet grasslands from the edges of the marshes, having a unique vocal repertoire and a unique plumage. Because of its restricted geographical distribution and the threats that have their habitat, this new species should be categorized as Vulnerable on the IUCN Red List.

## RESUMEN.-

Describimos una nueva especie de capuchino del género *Sporophila* (Emberizidae) de los Esteros del Iberá, provincia de Corrientes, en el noreste de Argentina. Esta especie habría permanecido sin identificar debido a la falta de exploraciones ornitológicas en la región central de los Esteros del Iberá y además habría sido confundida con individuos inmaduros de otras especies de *Sporophila*. Realizamos observaciones de comportamiento y hábitat, experimentos de playback, análisis comparativos de las vocalizaciones y comparaciones del plumaje con otras especies simpátricas del mismo género encontrando que esta especie, a la que nombramos como *Sporophila iberaensis*, habita pastizales húmedos de los bordes de los esteros, presentando un repertorio vocal y un plumaje únicos. Debido a su distribución restringida y las amenazas que tiene su hábitat, esta nueva especie debe ser categorizada como Vulnerable en la lista roja de la IUCN.

### SPOROPHILA

IS A Neotropical genus of ca 30 species of small (8-10 g) seedeaters of forests, grasslands and wetlands from southern United States to southern South America (Ridgely and Tudor 1989). In recent years a group of smaller species of *Sporophila* known as capuchinos, that inhabit southern South America grasslands, have attracted increasing interest by researchers who have studied their plumage, vocalizations, breeding ecology, systematic and evolution (Lijtmaer et al. 2004, Facchinetti et al. 2008, 2011, Areta 2008, 2009, Areta et al. 2011, Benites et al. 2010, Machado and Silveira 2011, Campagna et al. 2009, 2012, Areta and Repenning 2011a, 2011b, Repenning et al. 2010). In spite of diverse plumages, vocalizations, and habitat uses of the species of this group some studies indicate that they are genetically very similar (Lijtmaer et al. 2004, Campagna et al. 2009, 2012). This decoupling between the homogeneity of the genetic markers and phenotypic diversity would be best explained as a group that had undergone a recent radiation beginning in the Pleistocene, leaving genetic signatures of incomplete lineage sorting, introgressive hybridization and demographic expansions (Campagna et al. 2012). Another notable aspect of the capuchinos is their link with conservation issues. Capuchinos have a strong dependence to remaining natural grasslands and they have been used as indicator species for detecting important regions or sites for conservation such as Endemic Bird Areas (EBA, Stattersfield et al. 1998) and Important Bird Areas (IBAs, Devenish et al. 2009). Finally, most capuchinos are globally threatened because their small or geographically restricted populations are subject to habitat loss and capture for the illegal wildlife market (BirdLife International 2012) and because mainly mature colored males are targeted, this may have other demographic implications (Azpiroz et al. 2012).

In October 2001 and October 2002, A.S.D.G. studied the avifauna of grasslands in Esteros del Iberá and Aguapey River basin, in the province of Corrientes, NE Argentina (see Di Giacomo 2005 and Di Giacomo et al. 2010). During this fieldwork he recorded many individuals of the genus *Sporophila*. As noted in his field-notebook, on 18 October 2001 he observed two individuals of *Sporophila* in a wet grassland area 14 km S of Galarza (28° 12′ 55″ S 56° 43′ 32″ W) that were not identified as they exhibited plumages which seemed to be incomplete. These individuals had cream or dull colored underparts that looked more like females of the genus, but they had a conspicuous gray crown, black collar, brown wings, back and rump and black bill. These birds were observed together with other individuals of *S. palustris* and *S. cinnamomea*.

Between 2007 and 2011, A.S.D.G. studied the birds of the grasslands in the Esteros del Iberá, in Corrientes Province, Argentina, as part of a long term project for evaluating changes in the grassland avifauna as a result of livestock grazing. During this fieldwork A.S.D.G. recorded many individuals from six species of capuchinos of the genus *Sporophila* every year.

In the annual surveys on November and December of 2008 and 2009 A.S.D.G. observed additional unusual individuals with same plumage as the unidentified seedeaters previously observed near Galarza in 2001. In 2008 these seedeaters were recorded as immature *S. ruficollis*. However, during 2009 these birds were noted as “new form” because they were often observed in pairs and had breeding territories, similar to other *Sporophila* that regularly breed in the region. Females were similar to the other species of the genus. Male songs were noted as different from recordings of voices of other sympatric species including those with a similar plumage (*S. ruficollis* or *S. cinnamomea*). Expanding the search of these birds in a vast area of the region, it was clear that these undescribed seedeaters were defending territories in a particular type of habitat and that they were probably nesting in sites where other *Sporophila* were rare or absent. During this survey distinctive songs of two individuals separated by 50 km (Estancia San Nicolás y Cambyretá Reserve) were recorded, and they were used to assess the playback response of other new individuals that were observed in the grasslands. A total of 15 sight records of this unknown seedeater (total of 19 individuals), were obtained in November and December 2009. In 8 of these encounters he was able to assess their playback response, obtaining positive responses to the recording of the unidentified *Sporophila* and negative for recordings of other sympatric species.

During November and December 2010, B.L.L., C.K. and A.S.D.G. obtained an additional 23 records (35 individuals) covering a larger number of sites. They observed pairs on breeding territories and conducted additional playback trials. Most individuals recorded were paired with males defending territories and singing in response to the same distinctive song of the unidentified *Sporophila* recorded in 2009. These seedeaters did not respond to the voices of the other sympatric species of the genus that had similar plumage characteristics (*S. cinnamomea* and *S. ruficollis*), and they did not respond to vocalizations of other species of the genus (*S. palustris*, *S. hypochroma*, *S. hypoxantha* and *S. pileata*). In February 2011, the authors returned to the field to continue the observations and recorded 11 records and observed 16 individuals. Some of the individuals were found in the same territories as in the November and December 2010 survey at the Estancia San Nicolás. A.S.D.G. and C.K. observed one pair feeding two young birds. Finally, at Estancia San Alonso, B.L.L. used playback to test the response, recorded more vocalizations, and then collected one individual for further study.

Authors recorded new observations (31 records, 39 individuals) and performed new playback trials during the annual survey to the Iberá grasslands in November and December 2011, and January 2012. Six observations occurred at the same point counts as in 2010, and another two observations occurred at the same point counts as in 2009. From a total of 14 playback trials in 2011, 13 individuals responded positively to the song of the unidentified *Sporophila*. These birds did not respond to the songs of other sympatric species, and only one individual did not respond to any stimulus. Two of the individuals essayed were sound-recorded and then collected near Estancia Don Luis and at Estancia El Tránsito. Additionally, B.L.L. obtained other 3 records (4 individuals) from the Aguapey River Basin, during a visit in November 2011.

After analyzing the information obtained from our 81 field records during our years of fieldwork in the area and our knowledge of seedeaters of the genus *Sporophila*, and our study of museum specimens, we found that both the plumage and vocalizations were consistently different from all other sympatric species of *Sporophila*. We have also observed that the main population of these unidentified *Sporophila* was quite common in the distinctive wet grassland habitat that is dominated by flooded *Paspalum* tall grasses surrounding marshes in the sandy hills area in the north of the Esteros del Iberá, locally called “lomadas arenosas” (sensu Carnevalli 1994). Thus, based on the differences of coloration of the plumage, vocalizations and reproductive habitat, we propose to name this new seedeater

> *Sporophila iberaensis*, sp. nov.
>
> Iberá’s Seedeater
>
> *Capuchino del Iberá* (Spanish)
>
> *Caboclinho-do-Iberá* (Portuguese)

### Holotype

Colección Nacional de Ornitología, Museo Argentino de Ciencias Naturales “Bernardino Rivadavia” (MACN), Buenos Aires, no. MACN-Or 72854. Skin, adult male, skull 100% ossified, testes 7 × 5 mm and 6 × 4 mm, netted in a wet grassland in Estancia San Alonso, Esteros del Iberá, department of Concepción, province of Corrientes (28° 18′ 10. 2″, 57° 26′ 25.5″ W). Collected 19 February 2011 by B.L.L., prepared by C.K. Sound-recorded by B.L.L.; recording archived at Macaulay Library, Cornell Laboratory of Ornithology, Ithaca, New York (accession numbers: MLNS to be confirmed). A copy of sound-record is archived in Colección Nacional de Ornitología, Museo Argentino de Ciencias Naturales “Bernardino Rivadavia” (MACN), Buenos Aires (accession numbers: to be confirmed) and available on Xeno-canto (www.xeno-canto.org) (accession numbers: to be confirmed).

### Diagnosis: Morphology

An emberizid, assignable to the genus *Sporophila* by the combination of bill shape, size and pattern of plumage coloration. In appearance is closely similar to sympatric members of the ‘capuchino group’ of the genus *Sporophila*. Males are totally separable in plumage from all other species of the genus. Females are similar to other capuchino species. Male differ in plumage from other capuchinos by contrasting forehead and crown lead gray with the nape and hindneck blackish brown, joining lores, ear-covers, chin, throat and neck (Fig. 1). The back and rump are brown or olive brown, and breast, abdomen, belly and undertail covers are pale yellow to cinnamon. All specimens show white at the base of the tail feathers and the white wing speculum (on the dorsal view of remiges) looks larger than other capuchinos. Compared with sympatric *S. ruficollis*, the new species has the blackish brown throat extending to the dorsal neck reaching to form a tight collar but does not extend towards the body beyond the throat, unlike *S. ruficollis* where dark color of the throat reaches the breast. Unlike the new species, *S. ruficollis* has lead gray in the crown extended to the dorsal neck and back, but the rump is rufous, similar to the underparts. It has recently been described a color morph of *S. ruficollis*, called ‘caraguata’ (Areta et al. 2011), but unlike *S. iberaensis*, the black collar is broader and extends ventrally to the breast, and the underparts, back and rump are rufous. Another species with which the new species shares some characteristics of plumage is *S. cinnamomea*, which in its full adult plumage has a rufous throughout the body, except the lead grey in the crown and brown wings and tail. *S. iberaensis* might look like a immature male of *S. ruficollis* or *S. cinnamomea* with incomplete plumage in which only the color of the crown (gray) and the throat or neck (rufous or blackish brown) coincides with adult males and the rest of the plumage is similar to a female. However, because other characteristics of morphology and maturation of plumage in capuchinos, together with their unique vocalizations, makes it possible to readily differentiate immature capuchinos from the new species (see Remarks).

**FIG. 1.**
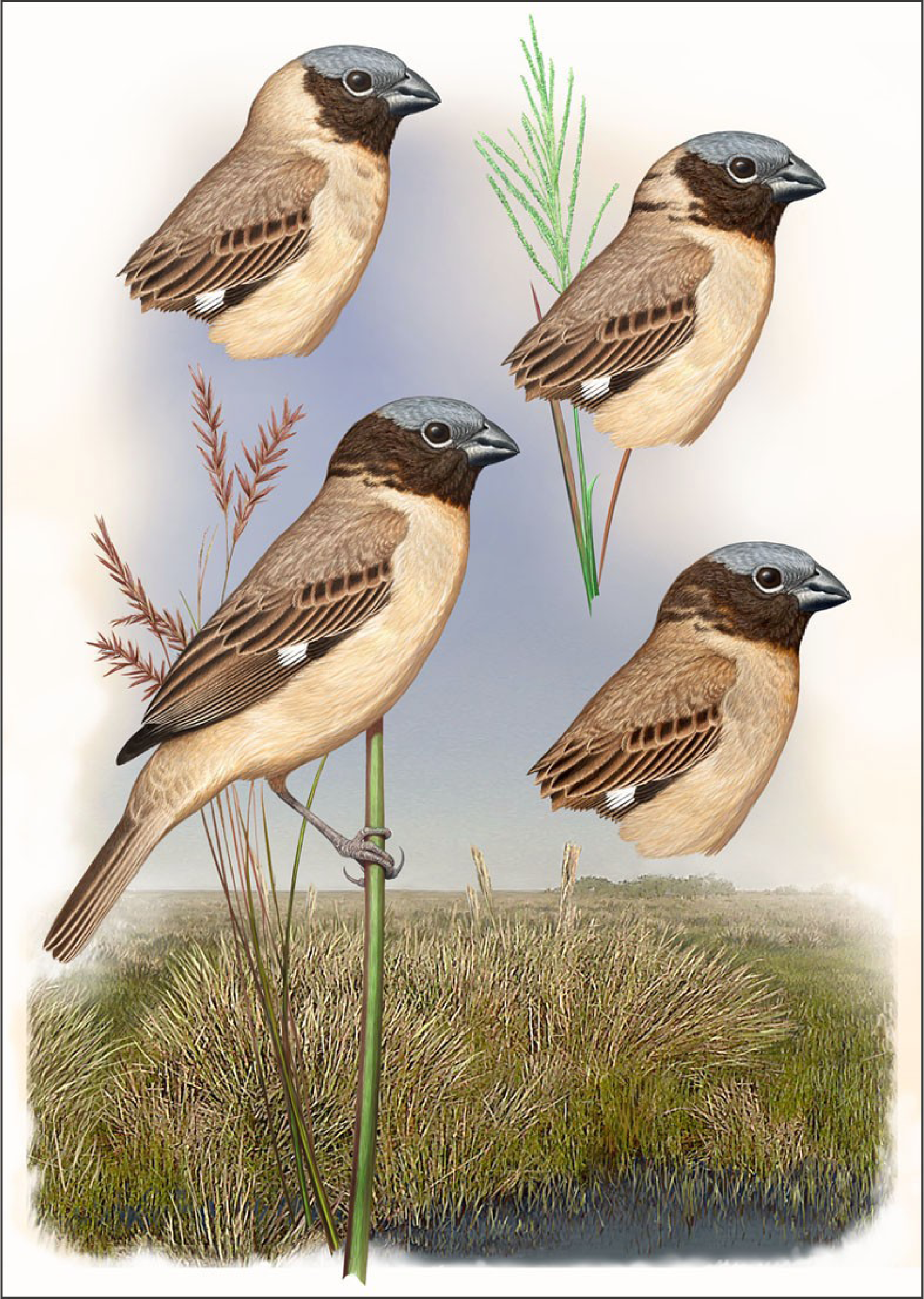
Plumage patterns of males Iberá’s Seedeater *Sporophila iberaensis* sp. nov. and their habitat in Esteros del Iberá, Corrientes, Argentina. Top left: An individual with incomplete collar. Top right: An individual with incipient dorsal collar and a detail of the grass *Paspalum rufum* whose seeds are eaten for seedeaters. Bottom left: An individual with full collar and fresh plumage at the beginning of the breeding season, and a detail of the grass *Andropogon lateralis*, the dominant plant of the Ibera’s grasslands. Bottom right: individual with full collar but with worn plumage as observed towards the end of the breeding season (based on holotype MACN-72854). Background: Typical habitat of *S. iberaensis* with wet grasslands dominated by *Andropogon lateralis* and *Paspalum* sp. around a marsh. Original painting by Aldo Chiappe.

Morphological measurements of *S. iberaensis* are in the range from those of other sympatric capuchinos of genus *Sporophila* (Table 1).

**TABLE 1.** Morphological measurements of *Sporophila cinnamomea*, *S. hypochroma*, *S. hypoxantha*, *S. iberaensis* sp. nov., *S. palustris, S. pileata* and *S. ruficollis* based on specimens listed in Appendix 3. Values are (in mm) mean ± SE with range and sample size between parentheses. Measurements of specimens were taken by A. S. D. G. following Baldwin et al. (1931). Bill length (exposed culmen) and tarsus were measured to the nearest 0.1 mm with calipers, and chord wing and tail length were measured with a metal ruler to the nearest 0.5 mm.

| Species | Bill length | Wing chord | Tail length | Tarsus length |
| --- | --- | --- | --- | --- |
| *S. cinnamomea* | 8.26±0.18 (7.8-8.7, n= 5) | 52.20±0.66 (50.0-54.0, n= 5) | 36.50±0.45 (35.0-37.0, n= 5) | 13.80±0.25 (13.0-14.5, n= 5) |
| *S. hypochroma* | 7.98±0.10 (7.4-8.2, n= 25) | 54.55±0.65 (51.0-58.0, n= 25) | 37.11±0.48 (35.0-40.0, n= 25) | 14.00±0.17 (13.5-15.0, n= 25) |
| *S. hypoxantha* | 8.17±0.07 (7.8-8.5, n= 30) | 53.82±0.55 (51.0-57.0, n= 30) | 37.30±0.37 (36.0-39.0, n= 30) | 13.72±0.19 (13.0-15.0, n= 30) |
| *S. iberaensis* | 8.1±0.0 (8.1-8.1, n= 3) | 52.33±0.88 (51.0-54.0, n= 3) | 40.5±0.5 (40.0-41.0, n= 2) | 13.33±0.33 (13.0-14.0, n= 3) |
| *S. palustris* | 8.30±0.26 (7.8-8.7, n= 3) | 55.25±0.25 (55.0-56.0, n= 4) | 36.50±0.29 (36.0-37.0, n= 4) | 12.83±0.44 (12.0-13.5, n= 4) |
| *S. pileata* | 8.54±0.09 (8.1-9.1, n= 14) | 53.97±0.31 (52.0-56.0, n= 15) | 36.93±0.44 (34.0-40.0, n= 15) | 13.70±0.18 (12.5-15.0, n= 15) |
| *S. ruficollis* | 8.12±0.07 (7.6-8.8, n= 54) | 54.62±0.25 (53.0-57.0, n= 52) | 37.53±0.34 (35.0-40.0, n= 53) | 13.56±0.19 (12.5-15.0, n= 51) |

### Diagnosis: Vocalizations

The vocalizations of *S. iberaensis* showed a typology common to the other capuchinos composed by introduction, main song and calls. However, each species of capuchino has certain characters in the main song or in the introduction, that are diagnostic (Narosky and Yzurieta 1987, Ridgely and Tudor 1989, Straneck 1990, Areta 2008, Areta et al. 2011, Areta and Repenning 2011a, Campagna et al. 2011). The vocalizations of *S. iberaensis* present two diagnostic notes that are very different from the homologous notes present in the vocalizations of other species of capuchinos. One note is present in the introduction, looking at the spectrogram with shape of an “Eiffel Tower” (Fig. 2A, C, D and E), reaching 5 to 7 kHz with greater amplitude (volume) in relation to other notes of the species. The other diagnostic note is present at the beginning of the main song (Fig. 2G, H, I, J, K and L), and compared to the homologous note in other species of capuchinos (Fig. 3), in *S. iberaensis* this note is unique because of its long duration, vibrato, “nasal” type of sound and notoriety of the harmonics (see detail in figure 2K). Both notes are very conspicuous to the human ear which makes it easy to differentiate this species from any other *Sporophila* in the field.

**FIG. 2.**
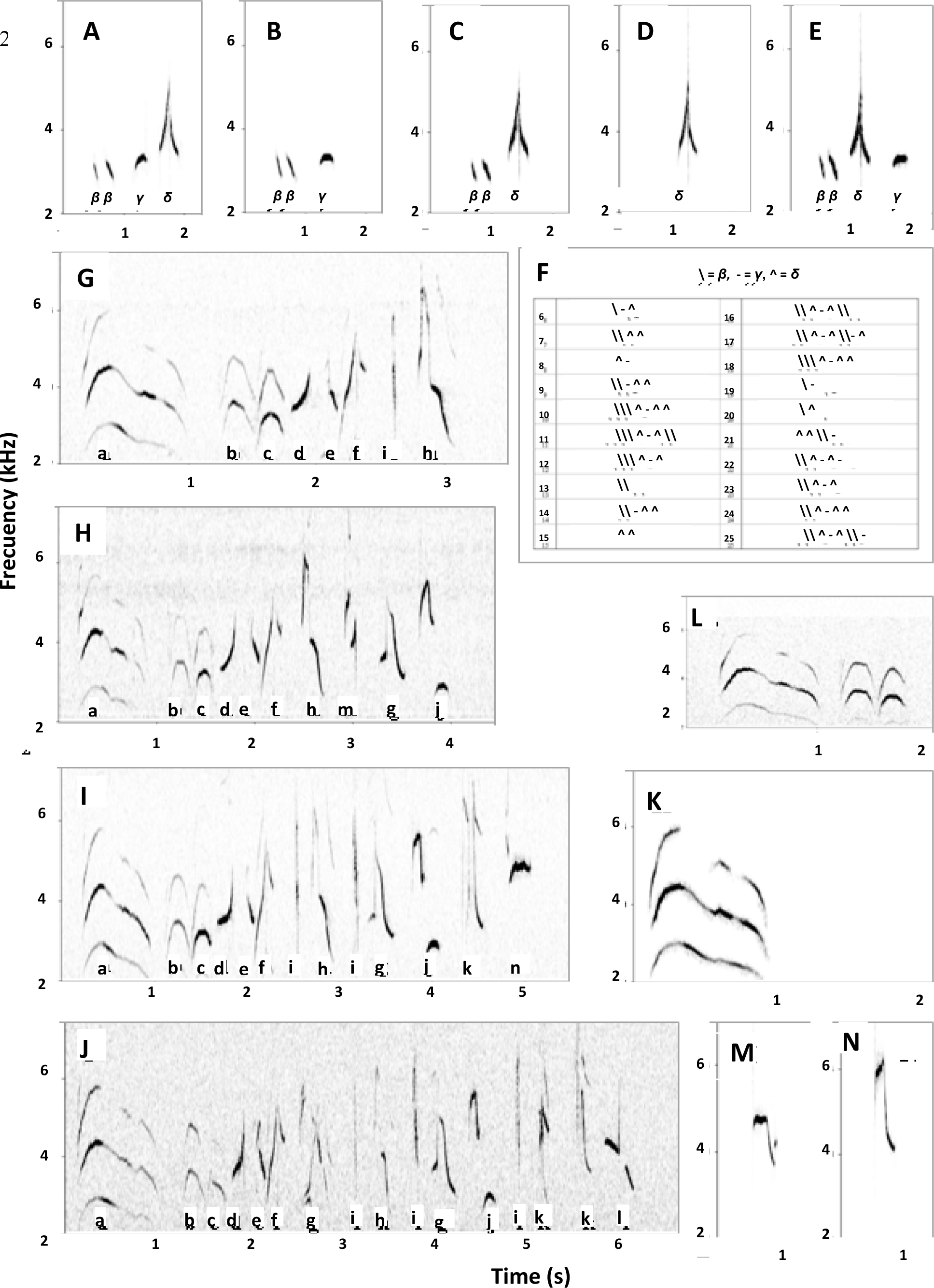
Representative audiospectrograms of vocalizations of *Sporophila iberaensis* sp. nov.: A-E, types of combination of notes (β, γ, δ) in introductions represented with more frequency in the samples. F, combination of notes in other introductions recorded less frequently. G-K, representative main songs and combination of notes (a, b, c, e, f, g h, i, j, k, l, m, n). L, detail of the principal note in the main songs. M-N, main calls. A, C, D, E, G, K, L and M, recorded by B.L.L. from holotype MACN-72854 (Estancia San Alonso, Concepción, Corrientes, Argentina, 19 February 2011). B and I, H, and J recorded by B.L.L. from other three different individuals (Estancia San Alonso, Concepción, Corrientes, Argentina, 27-28 December 2010 and 20 February 2011). K, recorded by A.S.D.G. (Cambyretá, Ituzaingó, 11 November 2009).

**FIG. 3.**
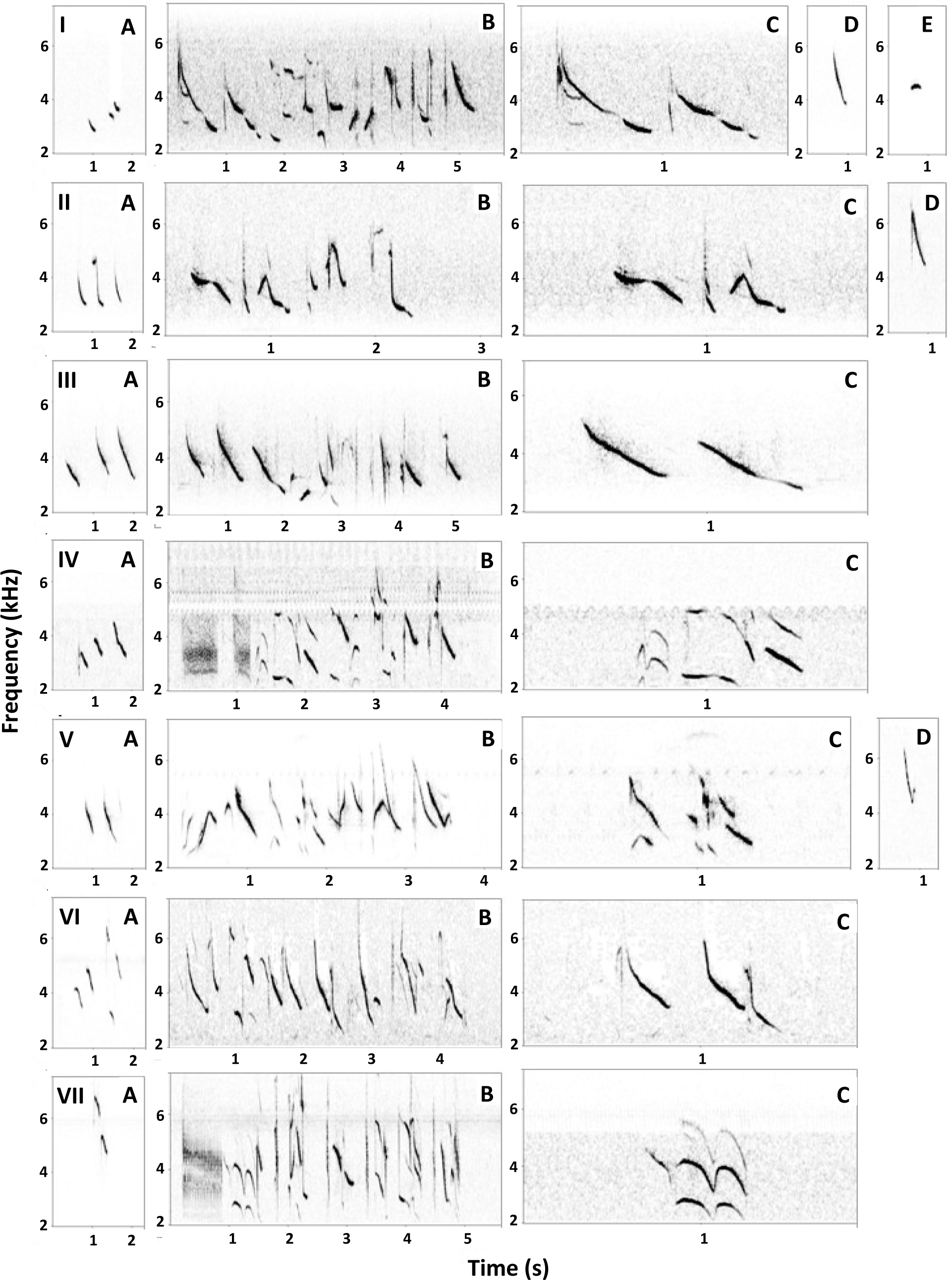
Representative audiospectrograms of vocalizations of other species of capuchinos (*Sporophila* sp.): I, *S. ruficollis*. II, *S. cinnamomea*. III, *S. hypoxantha*. IV, *S. hypochroma*. V, *S.palustris*. VI, *S. pileata*. VII, *S. melanogaster*. For each species is represented: A introduction, B main song, C detail of the diagnostic note in the main song, D main call (only for *S. ruficollis*, *S. cinnamomea* and *S. palustris* because they were identified as diagnostic by Areta 2008 and Areta et al. 2011). Records: I, *S. ruficollis* from Puente Barral, partido 25 de Mayo, Buenos Aires, Argentina, 4 February 2006 (López-Lanús 2008); II, *Sporophila cinnamomea* from Estancia Las Cruces (a, b,c), Gualeguaychú, Entre Ríos, Argentina, 2 December 2001 (López-Lanús 2008) and Ceibas (d), Entre Ríos, Argentina, 18 December 2000 (S. Wasylyk en López-Lanús 2008); III, *Sporophila hypoxantha* from Cruce al Puente Fray Bentos, Gualeguaychú, Entre Ríos, Argentina, 3 December 2001 (López-Lanús 2008); IV, *Sporophila hypochroma* from San Javier, Santa Fe, Argentina, 30 November 2004 (López-Lanús 2008); V, *Sporophila palustris* from Estancia El Potrero (a, b, c), Gualeguaychú, Entre Ríos, Argentina, 27 November 2008 (B. L. L.) and Estancia Santa Isabel (d), Alvear, Corrientes, Argentina, 4 December 2003 (R. Fraga en López-Lanús 2008); VI, *Sporophila pileata* from Estancia Virocay, Virasoro, Corrientes, Argentina, 15 December 2010 (B. L. L.); VII, *Sporophila melanogaster* from Rio Lava Tudo Lages, Santa Catarina, Brasil, 12 February 2008 (M. Repenning en López-Lanús 2008).

### Description of the holotype

See Figure 1 (individual at bottom right). Bill and legs black. Forehead and crown Medium Neutral Gray (capitalized color names follow Smithe 1975). Incomplete collar with lores, ear-covers, chin and throat Dusky Brown. Nape Olive Brown with some feathers Dusky Brown. Some underparts of the collar with feathers Cream Color. Back, scapulars, rump and upper tail coverts Olive-Brown. Breast, abdomen, belly and undertail coverts vary between Cinnamon and Cream Color. The side of the breast has a small spot in Dusky Brown. Upper wing coverts and alula Dusky Brown with borders in Olive-Brown. Primaries 9^th^ to 8^th^ Dusky Brown with anterior border in Olive-Brown. Primary 7^th^ Dusky Brown with anterior border Olive Brown and base of the posterior vane white. Primary 6^th^ to 2^nd^ Dusky Brown with anterior border Olive Brown and base of the posterior and anterior vane white. Primary 1st Dusky Brown with anterior border Olive Brown. Secondary 1st Dusky Brown with anterior border Olive Brown. Secondary 2^nd^ Dusky Brown with anterior border Olive Brown and base of the posterior vane white. Secondary 3^rd^ to 6^th^ Dusky Brown with anterior border Olive Brown and base of the posterior and anterior vane white. Primaries 8^th^ to 6^th^ with notch. Tertials Dusky Brown with borders Olive Brown. From the dorsal view the wings look Dusky Brown with a conspicuous white speculum that is broken by the absence of white between the Primary 1st and Secondary 1st. From a ventral view the wings look mainly white because of the extended white color of the primaries and also white in the axillaries feathers and wing lining region. The wrist region and underwing primary coverts are Dusky Brown. Rectrices are Dusky Brown with borders Olive Brown and the base of the anterior vanes are white.

### Measurements of holotype

Bill length (culmen exposed) 8.1 mm; wing length (chord) 51 mm; tail length 40 mm; tarsus length 13 mm; total length 102 mm; wingspan 165 mm; body mass 8.5 g.

### Designation of paratypes

MACN-72855, skin, adult male, skull 100% ossified, testes 8 × 5 and 8 × 5 mm, netted in a wet grassland in a road near Estancia Don Luis, Esteros del Iberá, department of Ituzaingó, province of Corrientes (27° 51′ 5.4″, 56° 53′ 38.4″ W). Collected 13 December 2011 by B.L.L., prepared by C.K. Sound-recorded by B.L.L.; recording archived at Macaulay Library, Cornell Laboratory of Ornithology, Ithaca, New York (accession numbers: MLNS to be confirmed). A copy of sound-record is archived in Colección Nacional de Ornitología, Museo Argentino de Ciencias Naturales “Bernardino Rivadavia” (MACN), Buenos Aires (accession numbers: to be confirmed)… and available on Xeno-canto (accession numbers: to be confirmed). MACN-72856, skin, adult male, skull 100% ossified, testes 8 × 5 and 8 × 5 mm, netted in a wet grassland in Estancia El Tránsito, Esteros del Iberá, department of Concepción, province of Corrientes (28° 28′ 2.4", 57° 42′ 22.2" W). Collected 15 December 2011 by A.S.D.G., B.L.L. and C.K., prepared by C.K. Sound-recorded by B.L.L.; recording archived at Macaulay Library, Cornell Laboratory of Ornithology, Ithaca, New York (accession numbers: MLNS to be confirmed). A copy of sound-record is archived in Colección Nacional de Ornitología, Museo Argentino de Ciencias Naturales “Bernardino Rivadavia” (MACN), Buenos Aires (accession numbers: to be confirmed) and available on Xeno-canto (accession numbers: to be confirmed).

### Variation in the type series

The type series includes three specimens, all males, that share the same diagnostic morphological and vocalization characteristics. The specimen MACN-72855 has a more complete collar lacking of Cream Color feathers on the ventral side alike of the holotype MACN-72854. The specimen MACN-72856 has some feathers Dusky Brown in the posterior zone of the gray crown and the incomplete collar has a few feathers of Cream Color, less than holotype MACN-72854. The tail feathers in specimens MACN-72855 and MACN-72856 have the bases of the anterior and posterior vane in white.

### Etymology

The specific epithet is a latinized adjectival form referring to the species main range at the ecosystem of the Esteros del Iberá (Iberá’s marshes) in the province of Corrientes, Argentina. “Iberá” is a term derivate from guaraní native language that means “glittering waters”. Esteros del Iberá is a complex mosaic of lagoons, rivers, wet grasslands and subtropical forests with an extension of 14,000 km^2^ and most of the area is protected as a Provincial Reserve, including private and government properties. By naming this species, we wish to draw attention for the conservation of the Esteros del Iberá as an remarkable reservoir of a rich cultural and natural diversity of our country.

## REMARKS

### Morphology

Males of *S. iberaensis* are similar in plumage making it quite easy to identify this taxon, despite the fact that some individuals vary in the extension of the collar (Fig. 1). Our study of the specimens collected and our field observations on *S. iberaensis* indicate that there is a range of individual variation in the dark collar, which in some individuals appears as incomplete and in others it looks completely full. Such variability in the extent or intensity of the coloration is reported in males of other species of *Sporophila* (see Stiles 1996, Machado and Silveira 2011). We noted similar variation in plumage among males of other species of capuchinos in the field sightings in Corrientes.

We compared specimens of *S. iberaensis* with 133 museum specimens (males) of capuchinos of genus *Sporophila* from collecting sites in the provinces of NE Argentina at Museo Argentino de Ciencias Naturales “Bernardino Rivadavia” (MACN, Buenos Aires; Appendix 3). We found differences between the extension of the black throat and differences in color of the throat and the ventral region of adult males of *S. ruficollis*. Similarly, we found a variation in the intensity of the ventral coloration in the series of specimens of adult males of *S. hypoxantha* and *S. hypochroma*, ranging from very light (cream and cinnamon) to very dark (rufous and marron). Such variability in intraspecific coloration means that most identifications in the field should be confirmed by their vocalizations.

Individuals of *S. iberaensis* were erroneously identified by A.S.D.G. as immature *S. ruficollis* during the field work prior to 2009. This is evident in the field notes of point counts; Table 2 compares the observed abundances of the capuchinos in a series of annual 100m-radious point counts of birds in the grasslands of Esteros del Iberá made between 2008 and 2011 (A.S.D.G.). The abundances of *S. ruficollis* for 2008 were overestimated due to the assignation of individuals with supposed immature appearance, but these actually belonged to *S. iberaensis*. Beginning in 2009, these birds were annotated as if they were another species on the basis of plumage and voice, and thus is evident that the annual counts had homogeneity in the rank of relative abundances of species between years 2009-2011.

**TABLE 2.** Annual variation in relative abundance (individual/point count), frequency (% of occurrence on total point counts) and rank order (order in relative importance of the species in the annual counts) of *Sporophila cinnamomea*, *S. hypochroma*, *S. hypoxantha*, *S. iberaensis* sp. nov.*, S. palustris, S. pileata* and *S. ruficollis* based on repeated 100m-radious points counts during the breeding seasons (November and December) of 2008, 2009, 2010 and 2011.

| Species | 2008<br>(n=115) | 2009<br>(n=126) | 2010<br>(n=104) | 2011<br>(n=83) |
| --- | --- | --- | --- | --- |
| *S. cinnamomea* | 0.08 (4%, 4) | 0.06 (2%, 4) | 0.08 (6%, 5) | 0.08 (6%, 4) |
| *S. hypochroma* | 0.03 (2%, 6) | 0.01 (1%, 7) | 0.15 (8%, 3) | 0.02 (2%, 5) |
| *S. hypoxantha* | 0.36 (17%, 1) | 0.09 (7%, 2) | 0.36 (21%, 1) | 0.17 (13%, 2) |
| *S. iberaensis* | - | 0.06 (5%, 3) | 0.27 (15%, 2) | 0.26 (15%, 1) |
| *S. palustris* | 0.04 (3%, 5) | 0.06 (3%, 5) | 0.06 (3%, 6) | 0.10 (6%, 3) |
| *S. pileata* | 0.10 (5%, 3) | 0.13 (5%, 1) | 0.15 (6%, 4) | 0.02 (2%, 6) |
| *S. ruficollis* | 0.15 (7%, 2) | 0.02 (1%, 6) | 0.06 (3%, 7) | 0.01 (1%, 7) |

Knowledge about the sequence of plumages in capuchinos is poorly documented in the literature; however it is important to compare available information regarding the differences between the maturation of the plumage of immature males of *S. ruficollis* and adult males *S. iberaensis*. An experiment carried out to study plumage molt of juvenile *S. ruficollis* extracted from their nests (1992 and 1993) and held in captivity for more than five years, found that immature males have blackish brown throats, but do not have a well defined the gray crown (M.A. Roda *in litt*.). During the first year, male *S. ruficollis* exhibit similar coloration than females, requiring more than a year to acquire their final male adult plumage. From 9 months of age, these young males have uneven spots, asymmetrical, of the blackish brown in throat, and chestnut in the rest of the ventral, both with different degrees of intensity in color. They acquired gray dorsal coloration at the end of maturation, when they have completed their full adult plumage, that is, when also have acquired much of the adult pattern and black-and-rufous color in the ventral region. Other character which allows differentiating immature birds in *Sporophila* is the horny coloration of the bill with pale or yellowish color in the basis, in adults the bill is uniformly black. Such characteristics of immature males of *S. ruficollis* are observed in museum specimens with age determination (Fig. 4).

**FIG. 4.**
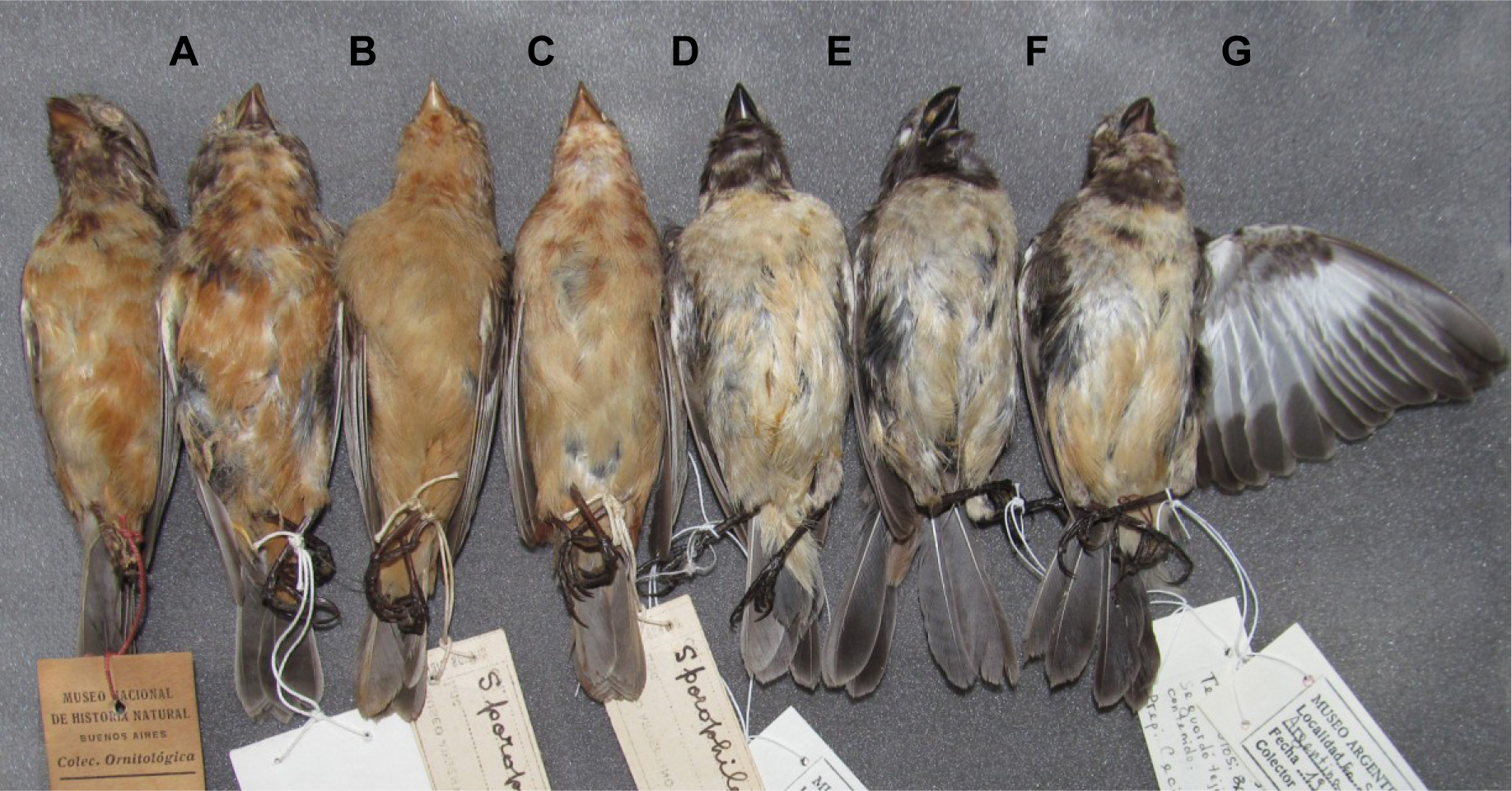
A-B, ventral view of immature males of *S. ruficollis* (MACN-Or-7436, MACN-Or-67324). C-D, ventral view of immature males of *S. cinnamomea* (MACN-Or-52377, MACN-Or-52376). E-G, ventral view of adult males of *S. iberaensis* sp. nov. (MACN-Or-72856, MACN-Or-72855, MACN-Or-72854). Immature individuals of sympatric capuchinos of genus *Sporophila* show the characteristic irregular pattern of mixed coloration on ventral side and notable yellowish bill.

These observations on plumage maturation of *S. ruficollis* agree with recent studies made with the species *S. hypoxantha* under laboratory conditions that were measured objectively using reflectance spectrometry (Facchinetti et al. 2011). Such study found that males have a plumage similar to females during their first year (delayed plumage maturation). Males acquire typical plumage of an adult after a year of age, being the crown the latest patch acquiring gray coloration. Further studies of the plumage of *S. hypoxantha* indicate that the pigments involved in reddish and dark coloration are melanins (transmission electron microscopy, Facchinetti 2012), while the olive or gray coloration of the crown and dorsal parts would correspond to structural (not pigment) and whose function is often associated with signals of male competitiveness or social dominance (see Facchinetti et al. 2011).

Study of museum aged specimens of *S. cinnamomea* showed similar characteristics than observed in bill and plumage of *S. ruficollis* (Fig. 4).

In three cases, it is considered that the *S. ruficollis*, *S. hypoxantha* and *S. cinnamomea*, exhibit delayed plumage maturation, a frequent phenomena in other species that similarly forage in mixed flock during the nonbreeding season and are sexually dichomatic (Lyon and Montgomerie 1986, Beauchamp 2003).

Finally, in relation to plumage maturation in *S. iberaensis* we only observed two solitary individuals with female-like plumage, which vocalized and behaved like males, perching high and responding strongly to playback.

### Vocalizations

As the voices of the capuchinos are species-specific and are diagnostic (Areta 2008, Areta et al. 2011, Areta and Repenning 2011a), we performed a comparative analysis of the characteristics of the vocalizations of *S. iberaensis* with other capuchinos species found in the province of Corrientes, Argentina. Knowing that *Sporophila* males exhibit strong territoriality during the breeding season, we also performed simple playback experiments in the field to assess the response of individuals *S. iberaensis* to different songs. Because individuals were not marked, we only considered for analysis data territorial males that were separated by more than 1 km.

For the comparative analysis of vocalizations we followed the methodology of Areta (2008) Areta et al. (2011) and Areta and Repenning (2011a, 2011b) because this method made it possible to resolve the taxonomic status of other previously recognized species and the identification of color morphs and voice dialects belonging to well-defined species. During our field work, we recorded vocalizations of 27 individuals of *S. iberaensis* using Zoom H4n and Edirol Roland Corporation recorders, with directional microphone Sennheiser ME-66 (copies of these sounds are deposited in MLNS, MACN and Xeno-Canto Archive, Appendix 4). We compared recordings of S. *iberaensis* with vocalizations of 101 individuals of seven capuchinos species (Appendix 4). We performed and examined spectrograms of the recordings using Syrinx 2.1 (John Burt, University of Washington). We characterized the notes that could be identified unambiguously in the songs of *S. iberaensis* on the basis of the distribution of the frequency and the relative position in the song despite variation among individuals (Fig. 2). We compared the introduction, main song and calls separately. We evaluated the occurrence of the each note in individuals of *S. iberaensis* and other capuchinos assuming they have the same general structure.

The introduction is composed of three basic types of notes (marked as β, γ and δ in Fig. 2A-E) that are combined alternating the order of appearance as shown in Figure 2F. We observed 25 different combinations in a total of 174 introductions and the five most frequent combinations representing 73% of all introductions are illustrated in Figure 2A-E. We did not find that any other species of capuchinos had the note shaped as an “Eiffel Tower” that is consistently present in the introductions of *S. iberaensis*. We consider this a diagnostic note of the species both considering our study of spectrograms of six species of capuchinos, and the spectrograms provided in the results of studies by Areta (2008), Areta et al. (2011), Areta and Repenning (2011a, 2011b) and Repenning et al. (2010).

For the main song of *S. iberaensis*, we identified 13 unambiguous notes, the first 6 notes (Fig. 2G-J, with marks a, b, c, d, e, f) were always emitted in the same order of all full songs we analyzed (N = 48 from 11 individuals). The other following notes that occur after (Fig. 2G-J, with marks g, h, i, j, k, l and m) were emitted by altering the order in some cases as shown in Figure 2. The diagnostic note (Fig. 2G-J with marks a, and 2K-L) is present at the beginning of the song. Compared to the homologous note in other species of capuchinos (Fig. 3I-VII, marked c), in *S. iberaensis* this note unique because of its long duration, vibrato, “nasal” type of sound and notoriety of the harmonics (Fig. 2K). The shape of this note was unchanged throughout all the songs we analyzed, and usually followed by two notes of similar form.

The two main calls *S. iberaensis* (Fig. 2M-N) are distinctive from other species such as *S. palustris*, *S. ruficollis*, *S. cinnamomea* and *S. hypoxantha* (Areta 2008, Areta et al. 2011, Areta and Repenning 2011a, 2011b) (Fig. 3). In *S. iberaensis*, the main calls are characterized by a stable frequency distribution and dense at the top of the notes before its fall vertically, independently of their frequency range. The first main call (Figure 2M) is characteristic by presenting at the end of the note a termination in a “V” small similar to the “nail” that bends downward that is described for the main call of *S. palustris* in Areta (2008) (Fig. 3VD). We consider the main call *S. iberaensis* diagnostic among other capuchinos.

The notes of the introduction, main song and calls of all other species of capuchinos are clearly different, both observed in our comparison, as with the direct comparison with the characteristic spectrograms for capuchinos that other authors have published (see Areta 2008, Repenning et al. 2010, Areta et al. 2011, Areta and Repenning 2011a, 2011b, Campagna et al. 2011). Note that in the introductions of other species (Figure 3I-VII), all notes have a frequency distribution almost vertical and descending in all cases, whereas in *S. iberaensis* these notes are ascending or stable, but mostly appears in the form of main note with shape of “Eiffel Tower” with a significant frequency range both ascending and descending symmetrically. In comparison to the main song of the other species of capuchinos, none were identical to the notes identified in *S. iberaensis* except for: 1) a note indicated by the letter i, which appears in the main song of *S. iberaensis* (Fig. 2I-J, marked with ‘i’), *S. cinnamomea*, *S. hypoxantha*, and *S. melanogaster* (Figs. 3IIB, 3IIIB and 3VIIB); 2) a note appears on the main song of *S. melanogaster* having a structure similar to diagnostic note of the main song of *S. iberaensis* when it appears repeated in “bc” position (not ‘a’ position), as show the spectrogram of Figure 2G-J vs Figure 3VIIC; 3) a note corresponding to the typical call *S. iberaensis* (Fig. 2N) resembles one of the notes of the introduction of S*. cinnamomea* (Fig. 3II, second note). In any case, the sequence of notes, the intensity of diagnostic notes and constancy of the voices of all the capuchinos analyzed is sufficient evidence of a distinctive vocal repertoire that is associated with individuals that have the characteristic plumage of *S. iberaensis*.

During the breeding seasons of 2009, 2010 and 2011 we conducted field observations to ensure that individuals characterized as *S. iberaensis* based on their plumage produced diagnostic vocalizations. We conducted playback experiments to 33 individuals with characteristic plumage of *S. iberaensis*. When one individual was singing we recorded their vocalization to obtain a good record. If the bird was calm and accustomed to the presence of the observer, we played songs of other capuchinos (*S. ruficollis*, *S. cinnamomea*, *S. palustris*, *S. hypoxantha*, *S. hypochroma*, and *S. pileata*), and then observed the reaction of the individual. If there was no response to the playback of another species, we completed the experiment playing the voice of an *S. iberaensis* recorded in 2009 and we assessed whether there was a response. We conducted this experiment to 22 individuals of *S. iberaensis* and we obtained negative results for the stimulus of the songs of another species, and positive response for the song of *S. iberaensis*. For 12 individuals, we played voices of the six species of capuchinos, varying the order of appearance, and for the other 10 individuals were reproduced voices of a single species (2 *S. palustris*, 5 *S. ruficollis*, 1 *S. hypochroma*, 1 *S. cinnamomea* y 1 *S. hypoxantha*). The positive response of the individuals to the voices of *S. iberaensis* was a sudden change in behavior, with quick changes of their perch, head movements, and strong approaches directly to the sound source. A further 11 individuals were tested first with the voices of *S. iberaensis* and responded positively. On one occasion an individual with plumage of *S. iberaensis* did not react to the song of *S. ruficollis*, nor to the sing of *S. iberaensis*.

We also note that our playing *S. iberaensis* vocalizations in their more characteristic habitats attracted individuals of this species on 3 separate occasions. We had not detected these birds previously, meaning that these individuals were attracted by the recordings from large distances (> 100 m), but were not attracted by such recordings any other species of capuchinos that are usually in these same habitats. Additionally, we performed other six experiments reproducing playback voices from *S. iberaensis* to other species of capuchinos in the area. We did not get positive responses to these recordings, but when we made playback of the “correct” species the response was positive.

### Distribution and habitat

During the fieldwork in the province of Corrientes in NE of Argentina, in 2001 and 2008-2011, we obtained a total of 81 records of *S. iberaensis* in 10 localities along Esteros de Iberá a quasi pristine region with a very low human population, as it has not suffered negative impacts from superficial inflow from water courses nor industrial activities in its borders (Fig. 5). Esteros del Iberá is one of the largest isolated inland freshwater wetlands located in South America, covering approximately14,000 km^2^. It is a flat ecosystem with a very gradual general slope with extreme altitude around 72 m a.s.l. at the northeast and 50 m a.s.l. to the southwest. The Iberá wetland consists of a vast mosaic of marshes, swamps and shallow lakes, most of which (60%) remain permanently inundated. Characteristic mats of floating vegetation called “embalsados” compose the shores and the surrounding environment of the lakes. The oriental region is characterized by a slow sheet flow where the biggest permanent shallow lakes are located in connection with the contiguous marshland. The central-western region, where *S. iberaensis* is more abundant, is characterized by an irregular topography which tends to form a plain and semi-fluvial drainage network of interconnected streams which collect the waters, and where emerging sandy hills conforming “islands” covered by grasslands and savannas (Fig. 6A). According to the flooding regime there are different types of habitats in Iberá which occur from the open water bodies to the “islands” where the terrain slightly rises and allows the development of terrestrial vegetation of grasslands and forests (Neiff 1981).

**FIG. 5.**
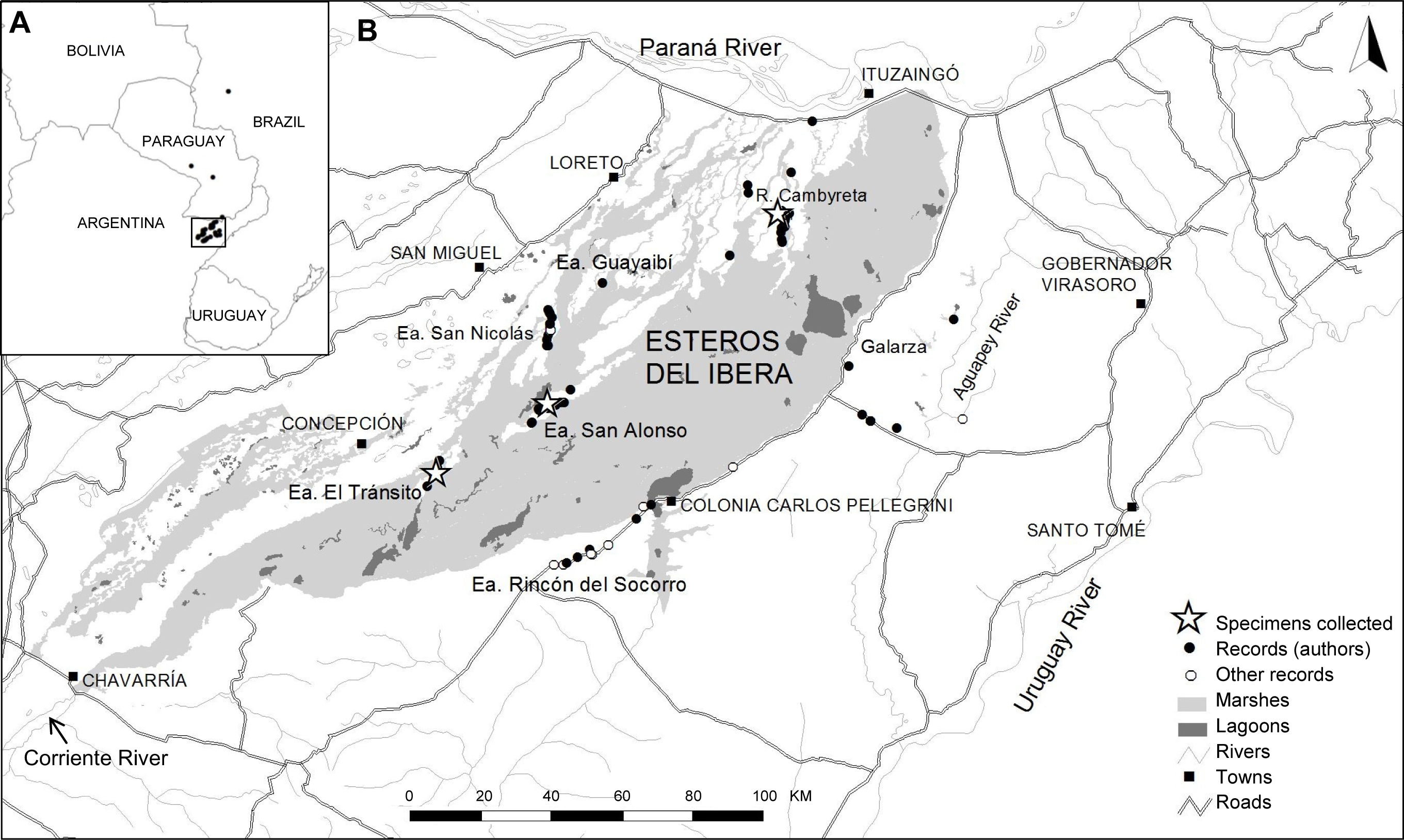
A) Map showing all known localities (black circles) for *Sporophila iberaensis* sp. nov. in Argentina, Paraguay and Brazil. B) Inset map of the province of Corrientes, Argentina, where are located the Esteros del Iberá showing the distribution of the records of *S. iberaensis* from authors (black circles) and other sources (white circles) around 10 localities. The stars represent the localities of the paratype series: MACN-Or-72854 (holotype, Estancia San Alonso), MACN-Or-72855 (Reserva Cambyretá), and MACN-Or-72856 (Estancia El Tránsito).

**FIG. 6.**
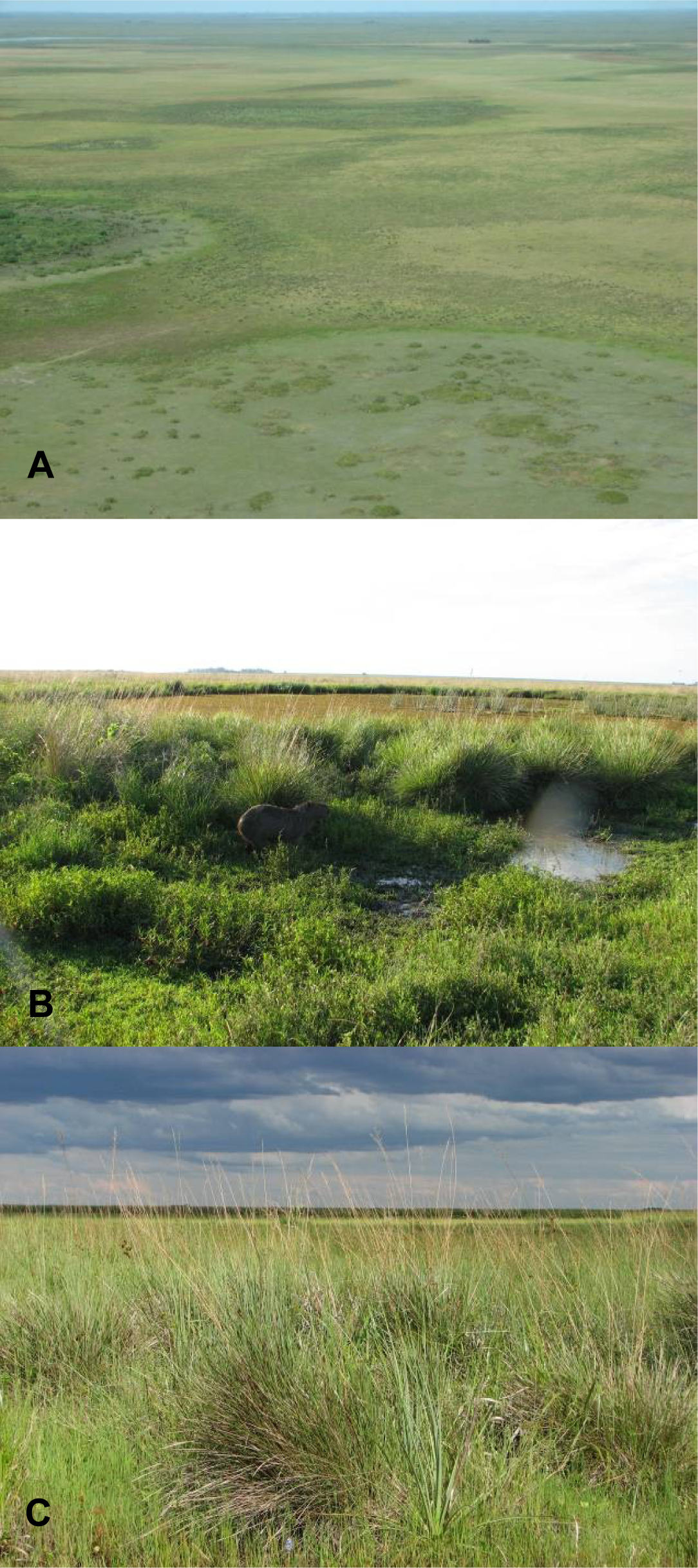
Landscape view of the Esteros del Iberá and habitat of *Sporophila iberaensis* sp. nov. A, Aerial photograph of sandy hills covered of wet grasslands of the Estancia San Alonso (type locality) along Paraná lagoon and Carambola stream (back of the image). B-C, Wet grasslands dominated by *Andropogon lateralis* and *Paspalum* sp. in territories of *S. iberaensis* in 2010-2011 (B, Estancia San Nicolás and C, near Estancia San Luis-Cambyretá). Photographs by A.S.D.G.

*S. iberaensis* inhabits wet tall-grasslands around vegetated marshes in the ecorregion of Esteros del Iberá. Such lowland grasslands are usually dominated by tall grasses such as *Paspalum durifolium* and *Andropogon lateralis* which produce seeds that are the main food of the capuchinos. These grasslands can be temporary flooded. A similar type of habitat is artificially created in Corrientes because of drainage channels or by retention of water along the roads. This is the reason that several records of *S. iberaensis* and another capuchinos are quite common along the roads. We found some territories of *S. iberaensis* very close to the water in the border of marshes with permanent water and covered of dense emerging aquatic plants of genus *Rhynchospora*, *Thalia*, *Panicum*, *Cyperus* and *Hymenachme*. Additionally B.L.L. observed three territorial males in the Aguapey River Basin during 2011, in an area of flood plains on sandy soil covered of grassland dominated by *Paspalum rufum*, similar to the physiognomy of the wet grasslands on sandy soils in the western Esteros del Iberá. No records of *S. iberaensis* are from more dry grasslands or savannas.

We obtained a set of additional 21 records through consultation with colleagues and bird photographers who had posted in Internet photos or recordings of *S. iberaensis* that were labeled as belonging to ‘young’ or ‘immature’ *S. ruficollis* (see Appendix 2). This set of photographic and voice records which may be assigned to *S. iberaensis* includes 15 records that fall exactly within of the 10 localities we studied in the province of Corrientes in Argentina, but it also includes six records that can expand the geographical distribution to three localities in Paraguay (San Cosme y Damián, department of Itapua; Bajo Chaco, department of Presidente Hayes; and Arroyo y Esteros, department of Cordillera) and one in Brazil (Aquidauana, state of Mato Grosso do Sul) (see Appendix 2, Fig. 5). For one photographic record obtained in Paraguay we did not achieve the exact location.

All localities for *S. iberaensis* are within the known geographical range of the group of southern capuchinos which occur sympatrically (see maps in Ridgely and Tudor 1989, Da Silva 1999, Campagna et al. 2011). However, in the case of *S. iberaensis*, to our knowledge, it seems to have a much more restricted geographical range in the province of Corrientes in Argentina, continuing northwards through the valley of the River Paraguay in Paraguay and Brazil.

Capuchinos are austral migratory birds that regularly breed in northern Argentina, Paraguay, Uruguay and southeastern Brasil, and migrate northwards in winter to central Brazil and Eastern Bolivia (Ridgely and Tudor 1989). Some capuchinos such as *S. hypoxantha* are partially migratory and maintain individuals during the winter in Formosa, in north Argentina (Di Giacomo 2005). To our knowledge records of *S. iberaensis* are from the breeding season with October 6 and March 11 as extreme dates, with no records during winter in Esteros del Iberá, thus the distributional range of this species during winter remain unknown.

### Ecology and behavior

*S. iberaensis* is a common species in its typical habitat in Esteros el Iberá. Like most of capuchinos that inhabit the region, we found *S. iberaensis* in wet or seasonally flooded grasslands that are dominated by large grasses with high vertical development and presence of many seeds in the spikes. We observed *S. iberaensis* eating the seeds of several grasses: *Andropogon lateralis*, *Paspalum durifolium*, *Paspalum rufum*, and *Hymenachme* sp. The seeds are consumed directly from the spikes; they do not eat the seed that fall on the ground. We observed *S. iberaensis* on two occasions eating fresh and green seeds from the spikes of *Hymenachme* sp. in February 2011 in Estancia San Nicolás, at the end of the summer when there are few seeds of *Paspalum* in the grasslands. One of the specimens of *S. iberaensis* that we collected in Estancia Alonso had green seeds of *Hymenachme* throughout its digestive tract.

During the breeding season males of capuchinos are usually on high perches of the grassland, and are very conspicuous and vocalizing much of the time. However, females are always hidden and fly lower, among the grasses. The males are very territorial, both to the intrusion of their territory by other capuchino of the same species or other species, and also to the playback of conspecific voices. In spite of high territoriality, in Corrientes it is common to find several species of capuchinos in the same site, even in the breeding season, where territories are contiguous but are mutually exclusive (A.S. Di Giacomo unpublished information). Thus, the fights are common among males defending contiguous territories.

We estimated the relative abundance of all sympatric capuchinos during the annual series of 100m-radious point counts of grassland birds in Esteros del Ibera. While the relative abundances of capuchinos varied between years, *S. iberaensis* always presented a high abundance relative to the other seedeaters (Table 2). Moreover, in some places of Estancia San Nicolás, Cambyretá and Carambola, in the region of sandy hills in central-western Esteros del Iberá, *S. iberaensis* seems to be the most abundant species.

We observed a pair of *S. iberaensis* feeding two fledglings in San Nicolás in February 2011. We also observed a female visiting a nest in a tall grass at the edge of a deep marsh in the territory of a male *S. iberaensis*, in November 2011. These data confirm that the species breeds in these habitats and during the same period as the other species of capuchinos. Additionally, the three males we collected showed active testes.

### Systematics and biogeography

Southern capuchinos show an extraordinary variation in phenotype (plumage and song) despite very low levels of genetic differentiation found in any of the genetic markers assayed (Lijtmaer et al. 2004, Campagna et al. 2009, Campagna et al. 2011). Similar to the Darwin’s Ground Finches (Freeland et al. 1999), the southern capuchinos could not be diagnosed using neutral markers despite extremely phenotypic differentiation. Recent studies on demographic expansions and gene flow between some of the species of southern capuchinos suggest a rapid, recent and ongoing continental radiation of the clade, with species diverging in male plumage coloration patterns and song (Campagna et al. 2011). According to the current practice of recognizing multiple species of southern capuchinos on the basis of plumage, song and reproductive habitat variation (Ridgely and Tudor 1989, Areta 2008, Areta et al. 2011, Areta and Repenning 2011a, 2011b, Campagna et al. 2011), *S. iberaensis* has a diagnosable plumage, song, and typical habitat for reproduction, including conspicuous territoriality of males with aggressive response to playback with conspecific song and ignoring the heterospecific song, which would support the existence of ongoing reproductive isolation.

Southern capuchinos are listed as ‘obligate’ grassland birds of the open habitats of South America, inhabiting savannas, grasslands and marshes in Cerrado, Beni, Pantanal, Chaco and Pampas ecosystems (Vickery et al. 1999), and reproductive areas of southern capuchinos are broadly overlapped with six species occurring in Esteros del Iberá, province of Corrientes, NE Argentina (Birdlife International 2012). Two other species of southern capuchinos do not inhabit in Argentina: *S. melanogaster* inhabits grasslands of altitude (700-1800 m a.s.l.) in coastal SE Brazil and *S. nigrorufa* restricted to savannas in SE Bolivia and Brazilian Pantanal. The reproductive area of *S. iberaensis* is centered in NE Argentina within the range of all currently recognized species of southern capuchinos, probably sharing the same evolutionary and biogeography history together all southern capuchinos, which require further studies to understand the decoupling between genetic and phenotypic traits involving new approaches such as field experiments on behavior and next-generation sequencing as suggested by Campagna et al. (2011).

Paleogeographically, Esteros del Iberá is a plain depression which constitutes the ancient bed of the Paraná River, and remained connected until the end of the Pleistocene when a geological uplifting of the eastern border of Esteros del Iberá occurred and the Paraná River moved towards the west to form the present riverbed (Neiff 1999). This displacement generated diverse wetlands from the floodplains associated to the displaced watercourses. From the end of Pleistocene the Iberá basin has been closed, mainly fed by rain and drains only towards the Middle Paraná through Corriente River in the south of the wetland (Fig. 5). In Esteros del Iberá, where *S. iberaensis* apparently has their unique breeding population, there are other endemic taxa such as a frog (*Argenteohyla siemersi pederseni*), two lizards (*Anisolepis longicauda* and *Liolaemus azarai*), three fishes (*Astyanax pynandi*, *Hyphessobrycon auca*, and *Trichomycterus johnsoni*) and six species of endemic plants (*Oxipetalum fontelae*, *Bernardia asplundii*, *Jatropa pedersenii*, *Portulaca meyeri*, *Picrosia cabreriana*, *Elatine lorentziana*) (Parera 2004). More recently, researchers have described an independent evolutionary lineage of tuco-tucos (small subterranean rodents of the genus *Ctenomys*) that inhabits sandy soils of the Esteros del Iberá (Caraballo et al. 2012, Gómez Fernández et al. 2012), in the same areas where was recorded *S. iberaensis*.

### Conservation

Two species of capuchinos of genus *Sporophila* are globally threatened (Birdlife International 2012): *S. palustris* (Endangered) and *S. cinnamomea* (Vulnerable), and two other are considered as Near Threatened (*S. ruficollis* and *S. hypochroma*). Main threats for capuchinos are habitat loss by conversion of natural grasslands to agriculture and forestation, and bird trapping. For such species the criteria to categorizing them as globally threatened is based on inferred or suspected population size reduction of ≥30% over the 10 years through decline of habitat or the level of exploitation, and decreasing population size under 10,000 mature individuals. All these globally threatened species have breeding populations in Corrientes, Argentina, mainly in Esteros del Iberá where there are seven Important Bird Areas identified under Birdlife International criteria (Di Giacomo 2005b). These species of threatened capuchinos also have breeding populations in other regions of Argentina (provinces of Entre Ríos, Santa Fe and Buenos Aires), SE Brazil (state of Rio Grande do Sul), SE and W Uruguay, SE Paraguay and a disjoint population of *S. hypochroma* is located E Bolivia (see Birdlife International 2012). To our knowledge the only breeding population of *Sporophila iberaensis* appears to be restricted to Esteros del Iberá and neighboring grasslands in Corrientes where the same threats exist for the other species of capuchinos. In terms of criteria for evaluating the conservation status of *Sporophila iberaensis*, the extent of occurrence in Corrientes can be estimated in less than 20,000 km^2^ and such subpopulation is estimating to contain no more than 1,000 mature individuals, which this species qualifies for the category of globally “Vulnerable” under the following IUCN Red List Criteria (IUCN 2001): A3cde+4cde; B1ab(i,ii,iii,iv); C2a(i). *S. palustris*, which has a much larger geographical distribution during the breeding season that *S. iberaensis*, including NE Argentina (provinces of Corrientes and Entre Rios), S. Paraguay, Uruguay and SE Brazil (Rio Grande do Sul) has been classified as “Endangered”. We suggest that until new population estimates are made for the capuchinos, it is important to categorize *S. iberaensis* at least as “Vulnerable”, both as a precaution as fit under the above IUCN criteria. Also, considering *S. iberaensis* as a threatened species that migrates to the grasslands of Paraguay and Brazil, it should also be included together other capuchinos in Appendix I of the Convention on Migratory Species where exist a Memorandum of Understanding (MoU) between Argentina, Brazil, Paraguay and Uruguay targeting Southern South American grassland birds (Azpiroz et al. 2012).

Most records of *S. iberaensis* in the province of Corrientes have been made within the Iberá Natural Reserve, a provincial reserve which has a core conservation area on public land complemented with a ring of private land. The public land area is mostly composed of large marshes, ponds and large seasonally flooded areas (about 700,000 hectares), which are not used for any productive activities. Surrounding this depression, the ring of private land areas is located in mainland and in the edges of the marshes that are occupied by large areas of grasslands devoted to cattle, afforestation and conservation (about 500,000 hectares). We recorded most *S. iberaensis* in the areas of mainland and edges of marshes dedicated to conservation (about 150,000 hectares belonging to an NGO The Conservation Land Trust). Today, private reserves bordering Iberá Natural Reserve represent a key area for the conservation of this species because it is the only known breeding area.

## ACKNOWLEDGMENTS

The present study was conducted under Bird Habitat Division of USFWS (NMBCA 5115 and 4544) and BBVA Foundation (Iberaqua – BIOCON-04-100/05 Project) sponsored research. A.S.D.G. and C.K. are Research Fellow of CONICET. A.S.D.G. work was also supported by CONICET (PIP-114-201101-00329) and C.K. was also supported by IDRC (iBOL, International Barcode of Life Project). We also thank Dirección de Parques y Reservas de la Provincia de Corrientes and The Conservation Land Trust for allowing us to conduct this study at Reserva Natural del Iberá, and Dirección de Recursos Naturales (Fauna) for authorizing study and capture permits (Santiago Faisal, Marcelo Beccaceci, Bernardo Holman and Jorge Silva Core). We are grateful for logistical support and companion during fieldwork to Alejandro Di Giacomo, Ariel Ocampo, Sofía Heinonen, Ignacio Jimenez Perez, Sebastian Cirignoli, Yamil Di Blanco, Karina Spoerring, Ricardo Quintana, Fernando Sosa, Carlos ‘Yuyito’ Figuerero, Anibal Parera, Pascual Perez, Malena Srur, Marisi Lopez, Adrián Galimberti, Alexander Keshel, Mauricio Losada, Miranda Collett, Familia Steed and Patricia Haynes. We thank Miguel Angel Roda and Miguel Ferro for providing unpublished results of the experiments on plumages of *Sporophila*, Alejandro Di Giacomo and Amalia Suarez for identifying plant species, Pablo Tubaro and Yolanda Davies for providing access and assistance to the MACN collection, and Peter Vickery for reviewing and editing this manuscript. As admirable as a person and as an artist, we want to specially thank Aldo Chiappe his beautiful painting of the new capuchino species. This paper is dedicated to the memory of Don José Antonio Ansola (“Che Patrón”) one of the last authentic ‘gauchos’ of the Esteros del Iberá who managed his lands in a traditional way, keeping all the native culture and rich biodiversity intact, so that we can enjoy it in the future. Also is dedicated to the mentors of The Conservation Land Trust, Douglas and Kristine Tompkins, who have established these same lands in strict conservation reserves and are working to recover populations of the most endangered species of the region.

## APPENDIX 1

Details of records of *S. iberaensis* sp. nov. obtained by authors (A. S. D. G., B. L. L. and C. K.) during this study in the province of Corrientes, Argentina. Records are grouped in localities as shown in Figure 5 (black circles in the inset map).

Ituzaingó (1). One record on the National Road N12 13 km of access to Ituzaingó (27° 38′ 3.6″ S 56° 48′ 48.4″ W), 10 November 2009, photo and voice recorded, A. S. D. G. and Ariel Ocampo.

Reserva Cambyretá (2): Three records in the reserve (27° 45′ 18.9″ S 56° 51′ 44″ W, 27° 51′ 12.6″ S 56° 52′ 3.5″ W, and 27° 54′ 56.9″ S 56° 53′ 5.5″ W) on 11 November 2009, voice recorded, A. S. D. G. and Ariel Ocampo; two records (27° 52′ 14.7″ S 56° 52′ 41.1″ W and 27° 57′ 9.2″ S 57° 0’ 29.9″ W) on 12 November 2009, A. S. D. G. and Ariel Ocampo; six records (27° 51′ 7″ S 56° 53′ 37.1″ W, 27° 50′ 47.6″ S 56° 53′ 7.1″ W, 27° 51′ 26.6″ S 56° 54′ 8. 5″ W, 27° 53′ 15.6″ S 56° 53′ 7.5″ W, 27° 53′ 52.9″ S 56° 53′ 13.3″ W, and 27° 55′ 16.6″ S 56° 53′ 0.3″ W), on 16 December 2010, A. S. D. G.; four records in Estancia Don Luis (27° 50′ 51.7″ S 56° 53′ 5.4″ W, 27° 51′ 5.4″ S 56° 53′ 38.6″ W, 27° 51′ 7.8″ S 56° 53′ 47″ W, and 27° 51′ 9.7″ S 56° 53′ 34.8″ W), on 13 December 2011, voices recorded and one specimen collected, A. S. D. G. and B. L. L.; two records in Estancia San Ignacio (27° 53′ 3.1″ S 56° 52′ 41.3″ W and 27° 51′ 55.8″ S 56° 52′ 33.8″ W), on 13 December 2011, B. L. L.; two records in Estancia Yaguareté Corá (27° 48′ 15.9″ S 56° 57′ 54″ W and 27° 47′ 7.5″ S 56° 57′ 59.1″ W), on 13 December 2011, B. L. L.; five records in the reserve (27° 52′ 47.2″ S 56° 52′ 46.5″ W, 27° 51′ 7″ S 56° 53′ 37.1″ W, 27° 50′ 47.6″ S 56° 53′ 7.1″ W, 27° 52′ 14.7″ S 56° 52′ 41.1″ W and 27° 53′ 15.6″ S 56° 53′ 7.5″ W), on 13 December 2011.

Estancia Guayaibí (3): One record (28° 1’ 4″ S 57° 18′ 38″ W) on 30 December 2010, B. L. L.

Estancia San Nicolás (4): One record (28° 6’ 1″ S 57° 25′ 50.8″ W) on 8 November 2009, photo and voice recorded, A. S. D. G. and Ariel Ocampo; four records (28° 7’ 27.4″ S 57° 26′ 7.1″ W, 28° 6’ 54.8″ S 57° 26′ 9.8″ W, 28° 8’ 36.8″ S 57° 26′ 25.9″ W and 28° 9’ 12.4″ S 57° 26′ 34.2″ W), on 13 December 2010, A. S. D. G.; two records (28° 8’ 0.6″ S 57° 26′ 6.2″ W and 28° 10′ 0.9″ S 57° 26′ 32.2″ W) on 14 December 2010, A. S. D. G.; three records (28° 4’ 54″ W 57° 26′ 24.7″ S, 28° 10′ 1″ S 57° 26′ 23.8″ W, and 28° 6’ 54.8″ S 57° 26′ 9.8″ W) on 12 December 2011, A.S. D. G.; two records (28° 5’ 15.6″ S 57° 26′ 11.3″ S and 28° 5’ 28.3″ S 57° 26′ 8.9″ W) on 12 February 2011, A. S. D. G. and C. K.; one record (28° 6’ 54.8″ S 57° 26′ 9.8″ W) on 13 February 2011, A. S. D. G. and C. K.

Estancia San Alonso (5): two records (28° 16′ 18.6″ S 57° 23′ 13.5″ W and 28° 21′ 5″ S 57° 28′ 44.5″ W) on 12 December 2009, Ariel Ocampo; six records (28° 18′ 11.6″ S 57° 26′ 31.3″ W, 28° 18′ 29″ S 57° 25′ 20.8″ W, 28° 18′ 7.4″ S 57° 24′ 8.1″ W, 28° 18′ 10.1″ S 57° 26′ 22.5″ W, 28° 18′ 12.2″ S 57° 26′ 15.5″ W and 28° 18′ 11″ S 57° 24′ 39.9″ W) on 27 December 2010, voice records, B. L. L.; three records (28° 18′ 52.9″ S 57° 27′ 17″ W, 28° 19′ 16.2″ S 57° 27′ 41.1″ W, and 28° 19′ 5″ S 57° 27′ 44.9″ W) on 28 December 2010, voice records, B. L. L.; one record (28° 18′ 10.1″ S 57° 26′ 25.5″ W) on 19 February 2011, voice records, one specimen collected, B. L. L.; seven records (28° 18′ 11.8″ S 57° 26′ 14.7″ W, 28° 18′ 16.8″ S 57° 26′ 31.3″ W, 28° 18′ 44.7″ S 57° 26′ 59.7″ W, 28° 18′ 48.6″ S 57° 27′ 2.8″ W, 28° 19′ 0.2″ S 57° 27′ 1.7″ W, 28° 18′ 56.4″ S 57° 26′ 25.4″ W, and 28° 18′ 55.1″ S 57° 26′ 23.7″ W) on 20 February 2011, voice records, B. L. L.; two records (28° 18′ 11.6″ S 57° 26′ 31.3″ W, 28° 21′ 0.6″ S 57° 28′ 38.2″ W) on 14 January 2012, voice records, B. L. L. and Máximo López-Lanús; two records (28° 18′ 1.9″ S 57° 26′ 35.4″ W and 28° 18′ 20.4″ S 57° 25′ 41.9″ W) on 15 January 2012 voice records, B. L. L.

Estancia El Tránsito (6): one record (28° 26′ 25.1″ S 57° 41′ 54.3″ W) on 7 November 2009, A. S. D. G. and Ariel Ocampo; one record (28° 30′ 7.4″ S 57° 43′ 35.9″ W) on 8 November 2009, A. S. D. G. and Ariel Ocampo; one record (28° 28′ 18″ S 57° 42′ 46.6″ W) on 10 November 2010, A. S. D. G.; six records (28° 28′ 2.3″ S 57° 42′ 21.9″ W, 28° 40′ 11.7″ S 57° 22′ 11.9″ W, 28° 27′ 59.9″ S 57° 42′ 17.7″ W, 28° 26′ 25.1″ S 57° 41′ 54.3″ W, 28° 27′ 28.1″ S 57° 42′ 9.6″ W and 28° 27′ 59.9″ S 57° 42′ 17.7″ W) on 15 December 2011, voice record, A. S. D. G., B. L. L. and C. K.

Estancia Rincón del Socorro (7): two records (28° 34′ 41.8″ S 57° 13′ 50.1″ W and 28° 41′ 2.5″ S 57° 23′ 46.2″ W) on 12 November 2009, A. S. D. G. and Ariel Ocampo; one record (28° 39′ 8.7″ S 57° 20′ 28.2″ W) on 16 December 2011, A. S. D. G.

Colonia Carlos Pellegrini (8): one record (28° 32′ 45.8″ S 57° 11′ 44.9″ W) on 12 November 2009, A. S. D. G. and Ariel Ocampo.

Aguapey River (9): one record on the secondary road (28° 6’ 15.6″ S 56° 28′ 39.5″ W) 12 November 2009, A. S. D. G. and Ariel Ocampo; one record in Estancia La Sirena (28° 21′ 45.7″ S 56° 36′ 45.6″ W) on 26 November 2011, B. L. L.; three records in Estancia La Clarita (28° 19′ 54.5″ S 56° 41′ 41″ W, 28° 20′ 48.8″ S 56° 40′ 28″ W and 28° 20′ 48.8″ S 56° 40′ 28″ W) on 30 November 2011, voice records, B. L. L. and Adrián Galimberti.

Galarza (10): one record (28° 12′ 55″ S 56° 43′ 32″ W) on 18 October 2001, A. S. D. G.

## APPENDIX 2

Details of additional records of confirmed (available photograph and voice records) and possible (available photograph) *S. iberaensis* sp. nov. obtained through consultation with colleagues and bird photographers who had posted in Internet photos or recordings labeled as belonging to ‘young’ or ‘immature’ *S. ruficollis*. Records are grouped by localities as shown in Figure 5 (black circles in regional small map A, and white circles in the inset map B).

*Argentina, province of Corrientes*. Estancia San Nicolás (4): five records (28° 7’ 52.7″ 57° 26′ 0.6″) on 5 December 2011, voice record, Carlos Figuerero (*in litt.*). Estancia Rincón del Socorro (7): one record (approximated coordinates 28° 39′ 48.9″ 57° 20′ 6″) on 3 November 2008, one record on 17 October 2009, two records on 14 October 2009 and one record on 8 December 2009, Carlos Figuerero (*in litt*.); one record (28° 39′ 43.9″ 57° 20′ 16.5″) on 14 October 2010, Rosana Ursino and Pablo Mosto (<u>http://www.foto-mundosilvestre.com/data/media/1/capuchno_garganta_caf_0082.jpg</u>); one record (28° 41′ 19.7″ 57° 25′ 34.2″) on 14 January 2012, Ángel Luis Prato (<u>http://www.ecoregistros.com.ar/site/imagen.php?id=11257</u>). Colonia Carlos Pellegrini (8): one record in Cambá Trapo (28° 27′ 20.6″ 57° 0’ 1.4″) on 14 October 2009, Carlos Figuerero (*in litt.),* one record near the town (28° 41′ 15.5″ 57° 24′ 13.6″), on 11 March 2010, Natalia Villanova (<u>http://www.foto-mundosilvestre.com/data/media/10/juvenil_g.cafe.jpg</u>); Aguapey River (9): one record in Estancia La Clarita (28° 20′ 48.8″ 56° 40′ 28″) on 30 November 2010, Adrián Galimberti (*in litt*.).

*Brazil.* One record from Pousada Aguapé in Aquidauana, state of Mato Grosso do Sul (20° 5’ 44″ 55° 57′ 54″) on 9 January 2005, Josep del Hoyo (Internet Bird Collection, <u>http://ibc.lynxeds.com/species/dark-throated-seedeater-sporophila-ruficollis</u>).

*Paraguay.* One record from Arroyo y Esteros km 74, department Cordillera (25° 1’ 38.3″ 56° 51′ 32.6″) on October 2008, Paul Smith (Aves Paraguay, <u>http://www.faunaparaguay.com/sporophila_ruficollis.html</u>). One record in “Bajo Chaco” department Presidente Hayes (24° 23′ 3.5″ 58° 6’ 50.5″) from October 2008, Clyde Morris (Aves Paraguay, <u>http://www.faunaparaguay.com/sporophila_ruficollis.html</u>). Two records from San Cosme y Damian, department Itapua (27° 19′ 13.1″ 56° 19′ 53.6″) on 6 October 2011, Claudio Dias (Wiki Aves Brasil, <u>http://www.wikiaves.com/535628</u> and 469585). One record (not mapped) without date and locality from “Paraguay”, Frank Fragano (Aves Paraguay, <u>http://www.faunaparaguay.com/sporophila_ruficollis.html</u>).

## APPENDIX 3

List of specimens of genus *Sporophila* examined and measured for this study at Museo Argentino de Ciencias Naturales “Bernardino Rivadavia” (MACN, Buenos Aires, Argentina).

*S. iberaensis* (N=3): MACN-Or-72854, 72855, 72856.

*S. ruficollis* (N=53): MACN-Or-66435, 66054, 66130, 67354, 66075, 66076, 56172, 66131, 56118, 66011, 67335, 66913, 66243, 66053, 66012, 57955, 22430, 22424, 8560, 22423, 22425, 22419, 18563, 18567, 22422, 7436a/b/c/d/e, 18584, 22426, 22421, 22427, 18566, 6292a, 35183, 7349, 3791a, 8560a/b, 52745, 46176, 29992, 45375, 45374, 7349, 6293a, 29991, 45376, 31392, 4215a. *S. ruficollis ‘*caraguata’(N=1): MACN-Or-40039.

*S. hypoxantha* (N=30): MACN-Or-66966, 58606, 64295, 60348, 56166, 64260, , 58607, 60080, 58610, 56365, 63998, 68358, 66167, 67451, 66126, , 7190, 7235a/b, 7349a/b, 45378, 45370, 7436a/b, 29639, 29640, 45363, 45384, 45381, 45385.

*S. hypochroma* (N=25): MACN-Or-63967, 66894,66125, 66609, 66715, 66186, 59949, 66132, 66918, 48247, 45380, 22312, 22364, 40038, 40037, 48470, 45379, 22309, 22311, 40031, 43296, 45382, 43302, 40036, 68727.

*S. pileata* (N=15): MACN-Or-58605, 54925, 58604, 60774, 60773, 48220, 48234, 9002, 38164, 8560, 48237, 9002, 8560, 48243, 48222.

*S. palustris* (N=4). MACN-Or-59947, 8961, 48513, 52379.

*S. cinnamomea* (N=5). MACN-Or-52373, 52374, 52375, 52376, 52379.

## APPENDIX 4

List of recordings of capuchinos of genus *Sporophila* used for this study. Codes of the recordings correspond to the following collections: MACN, Museo Argentino de Ciencias Naturales “Bernardino Rivadavia”, Buenos Aires, Argentina); XC, Xeno-canto (www.xeno-canto.org) and BLL, private collection of Bernabé López-Lanús.

*S. iberaensis* (N=27): MACN-Sn 00001/2013 to 00041/43. XC and Cornell ‐TBC

*S. ruficollis* (N=28): XC47168, XC47170, XC49699, XC49700, XC49703, XC49701, XC51910, XC51911, XC51912, XC51913, XC51914, XC51915, XC51916, XC51917, XC51918, XC51919, XC51920, XC51921, XC108337, XC50171, XC50172, XC108334, XC108338, XC108335, XC108336, XC108339, XC108341, XC108342.

*S. hypoxantha* (N=13): XC51949, XC51950, XC51951, XC51952, XC51953, XC61760, XC108328, XC108327, XC108323, XC108324, XC108325, XC108326, XC108329.

*S. hypochroma* (N=18): XC45246, XC51937, XC51939, XC51940, XC51941, XC51942, XC51943, XC51944, XC51945, XC51946, XC61759, XC108322, XC108313, XC108314, XC108316, XC108318, XC108321, XC108320.

*S. cinnamomea* (N=11): XC65741, XC47162, XC47163, XC49694, XC49695, XC49696, XC49697, XC51908, XC61758, XC49698, XC108306.

*S. palustris* (N=19): XC17868, XC17997, XC42440, XC47161, XC84742, XC49707, XC49708, XC49709, XC49710, XC49712, XC51931, XC51932, XC51933, XC51934, XC94481, XC49711, XC108331, XC108333, XC108330.

*S. pileata* (N=10): XC49693, XC49692, XC15927, BLL293/23, BLL293/24, BLL293/25, BLL294/125, and three recordings from M. Castelino published in López-Lanús (2008).

*S. melanogaster* (N=2): XC13513, XC41808.

**FIG. 1.**
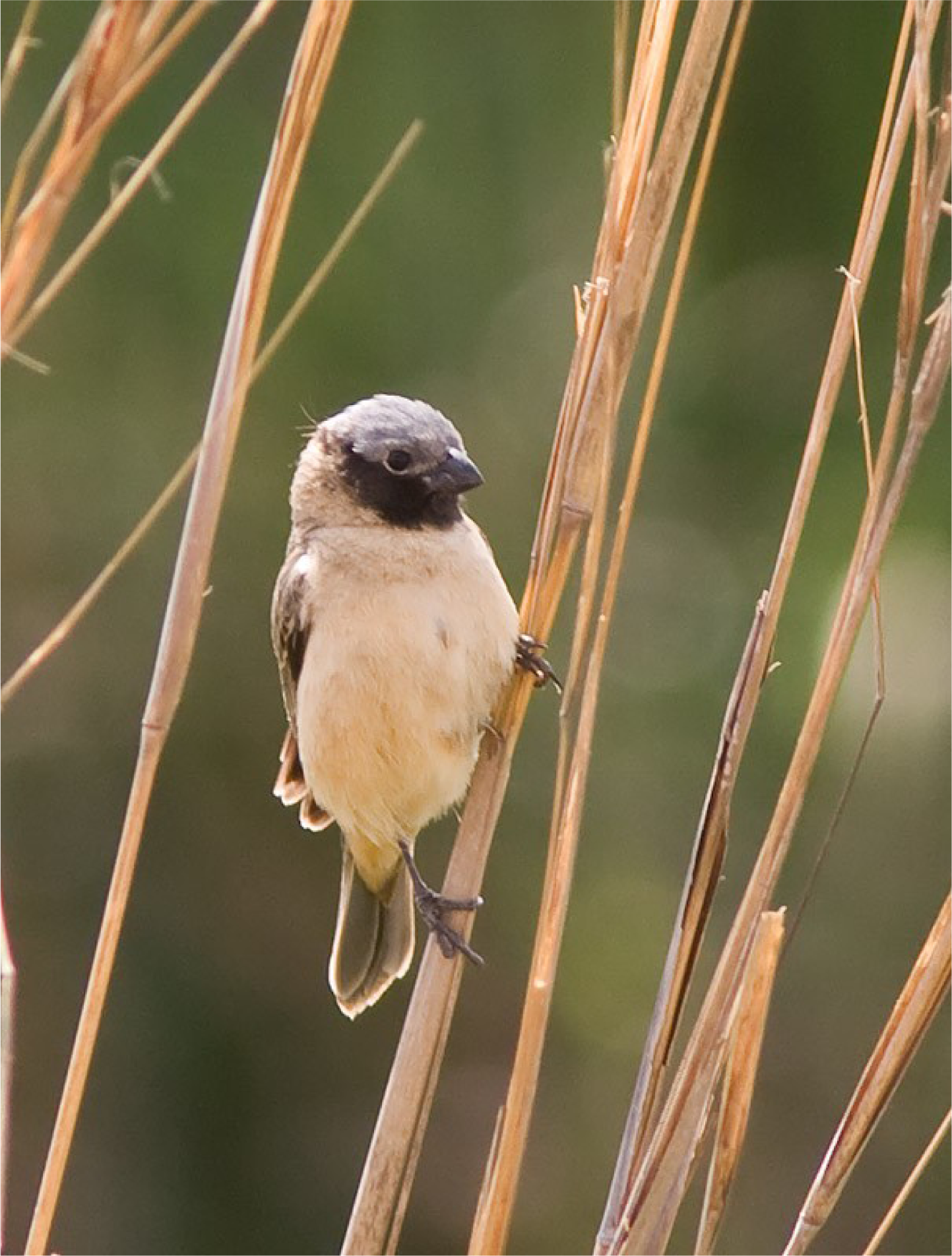
Optional (photo). Author: Carlos Figuerero.

